# N-Acetylcysteine alters disease progression and increases Janus Kinase mutation frequency in a mouse model of precursor B cell acute lymphoblastic leukemia

**DOI:** 10.1101/2023.10.19.563162

**Authors:** Mia P. Sams, James Iansavitchous, Madeline Astridge, Heidi Rysan, Li S. Xu, Bruno Rodrigues de Oliveira, Rodney P. DeKoter

## Abstract

B cell acute lymphoblastic leukemia (B-ALL) is the most prevalent type of cancer in young children and is associated with high levels of reactive oxygen species (ROS). The antioxidant N-acetylcysteine (NAC) was tested for its ability to alter disease progression in a mouse model of B-ALL. Mb1-CreΔPB mice have deletions in genes encoding PU.1 and Spi-B in B cells and develop B-ALL at 100% incidence. Treatment of Mb1-CreΔPB mice with NAC in drinking water significantly reduced the frequency of CD19^+^ pre-B ALL cells infiltrating the thymus at 11 weeks of age. However, treatment with NAC did not reduce leukemia progression or increase survival by median 16 weeks of age. NAC significantly altered gene expression in leukemias in treated mice. Mice treated with NAC had increased frequencies of activating mutations in genes encoding Janus Kinases 1 and 3. In particular, frequencies of *Jak3* R653H mutations were increased in mice treated with NAC compared to control drinking water. NAC opposed oxidization of PTEN protein ROS in cultured leukemia cells. These results show that NAC alters leukemia progression in this mouse model, ultimately selecting for leukemias with high *Jak3* R653H mutation frequencies.

## 1. Introduction

Precursor B cell acute lymphoblastic leukemia (pre-B-ALL) is the most frequently occurring type of cancer in young children. Despite a high remission rate, B-ALL is still the leading cause of cancer-related deaths in children, and more work needs to be done to reduce the long-term effects of chemotherapy toxicity [1]. Eighty per cent of pediatric ALLs are cancers of the B lymphocyte lineage (B-ALL), with 60% of these involving mutation or chromosomal translocation of genes encoding transcription factors [2]. Development of B-ALL is caused by initiating genetic lesions in developing progenitor/precursor B cells that function as drivers for the accumulation of secondary driver mutations, leading to oncogenic transformation by an evolutionary selection process [3,4]. Mutations that confer a survival or proliferation benefit are known as driver mutations, whereas passenger mutations do not provide a selection benefit [5,6]. Pre-B-ALL is thought to arise from large pre-B cells, a stage of development at which pre-B cell receptor and interleukin-7 (IL-7) signaling drive proliferation [7]. Downstream of IL-7 receptor signaling, Janus Kinases 1 and 3 (JAK1/3) activates Signal Transduction and Activator of Transcription-5 (STAT5) to translocate to the nucleus, and increase metabolism and reactive oxygen species (ROS) in leukemia cells [8].

Previous studies have shown that many types of human cancer, including leukemia, have higher ROS levels than in normal cells [9]. Lymphoid leukemias do not express pro-oxidant enzymes of the NOX family, so the origin of ROS is thought to originate from the electron transport chain [10]. Reactive oxygen species in the cell include superoxide, hydrogen peroxide, or hydroxyl radical [10]. Janus Kinase signaling has been implicated in the generation of high levels of ROS in leukemia cells [11]. High levels of ROS activate cellular signaling, including the Janus Kinase pathway, by diverse mechanisms including oxidation of protein tyrosine phosphatases, leading to activation of cellular metabolism [12]. However, the danger to cells from continuously elevated levels of ROS is oxidative damage, including 8-oxoguanine DNA damage that can lead to C->A transversion mutations [13,14]. Such mutations can contribute to clonal evolution of leukemias.

We have previously described a mouse model of B-ALL in which the related E26-transformation-specific (ETS) transcription factors PU.1 and Spi-B are deleted in B cells. This mouse model is termed Mb1-CreΔPB since conditional deletion of *Spi1* encoding PU.1 is under the control of the B cell-specific *Mb1 (Cd79a)* gene [15]. In these mice, there is a block in B cell development at the pre-B cell stage between the large pre-B and small pre-B steps resulting in a complete lack of mature B cells in tissues [15]. Consequently, this increases the large pre-B cell frequency and ultimately leads to the development of pre-B cell acute lymphoblastic leukemia (pre-B-ALL) at an incidence of 100% by a median 18 weeks of age [16]. Leukemia cells and cultured pre-B cells from Mb1-CreΔPB mice had high levels of ROS compared to wild type pre-B cells and had high levels of 8-oxoguanine DNA damage in their nuclei. Mb1-CreΔPB mice also had downregulation of most antioxidant genes, suggesting that these cells depend on high levels of ROS for survival and proliferation [17]. The addition of N-acetyl-L-cysteine (N-acetylcysteine, NAC) to the cell culture media inhibited the proliferation of cultured Mb1-CreΔPB leukemia cells and dramatically altered gene expression. NAC is the acetylated precursor of the amino acid L-cysteine. A substantial scientific literature shows that NAC functions as an antioxidant both in cell culture and *in vivo* by acting as a nucleophile for reactive oxygen species [18–20]. However, another mechanism for NAC is by serving as a precursor for synthesis of glutathione and in turn thiols, important antioxidants involved in many physiological processes [18]. The inhibition of proliferation of leukemia cells by NAC in culture suggested that proliferation requires ROS [17]. Therefore, we hypothesized that treatment with NAC *in vivo* would delay leukemia progression in the Mb1-CreΔPB mouse model.

To test this hypothesis, we compared the effects of NAC-containing or regular (control) drinking water provided to Mb1-CreΔPB mice starting at weaning. We found that treatment of Mb1-CreΔ1PB mice with 1.0 g/L of NAC in drinking water significantly reduced the frequency of CD19^+^ pre-B-ALL cells infiltrating the thymus at 11 weeks of age. However, treatment with NAC at 1.0 or 6.5 g/L did not reduce leukemia progression or increase survival by median 16 weeks of age. A concentration of 1.0 g/L NAC significantly altered gene expression in leukemias in treated mice. Mice treated with 1.0 g/L NAC had increased frequencies of activating mutations in genes encoding Janus Kinases 1 and 3. In particular, frequencies of *Jak3* R653H mutations were increased in mice treated with 1.0 g/L NAC compared to control drinking water. NAC could reduce *Phosphatase and tensin homolog deleted on chromosome ten* (PTEN) protein oxidized by ROS in cultured leukemia cells. These results show that NAC alters leukemia progression in this mouse model, ultimately selecting for leukemias with high *Jak3* R653H mutation frequencies. The implications of these findings for human disease are discussed.

## 2. Methods

### 2.1. Animal care

Mice were maintained in accord with guidelines of the Animal Care Committee at Western University and the Canadian Council on Animal Care. Mb1-CreΔPB mice were generated and maintained as previously described [15]. NAC-containing water was prepared using reverse osmosis filtered tap water and granulated N-acetyl-L-cysteine (>99%, MilliporeSigma Canada, Oakville ON) and was sterilized using 0.2 um PES filter units (VWR Canada, Mississauga ON). The prepared water was stored for a maximum of 2 months before use. Mice were weaned at 21 days of age, caged with 1-2 same-sex litter mates, and were randomly assigned to be administered either control water or NAC water. Water was administered through drinking bottles and was replaced with fresh water every 2-7 days. Water intake was measured by weighing bottles every 2-3 days.

### 2.2. Cell Culture

The preleukemic Interleukin-7-dependent pre-B cell line SeptMBr was established by culturing bone marrow from a 6-week old Mb1-CreΔPB mouse in Iscove’s Modified Dulbecco’s Media (IMDM, Wisent, St. Bruno Quebec) supplemented with 10% fetal bovine serum (Wisent), Penicillin-Streptomycin (Wisent), L-glutamine (Wisent), and 5% conditioned media from the Interleukin-7-producing cell line J558-IL-7 [21]. Continuously proliferating cells were passaged every 2-3 days and expressed CD19 at high levels as determined by flow cytometry.

### 2.3. Flow cytometry

Thymus was removed from euthanized mice, homogenized, and suspended in MACS buffer (500 mL 1x D-PBS, 1% 0.5M EDTA pH 8.0, 2.5 g BSA fraction V, MilliporeSigma). Single-cell suspensions were stained with Phycoerythrin-conjugated anti-CD19 antibody (clone 6D5, BioLegend, San Diego CA in preparation for flow cytometry. Live-dead discrimination was performed using Sytox Blue dye (Thermo-Fisher Scientific Canada, Markham ON). Flow cytometry was performed using a BD Cytoflex instrument (BD Biosciences, Franklin Lakes NJ).

### 2.4. RNA-sequencing

RNA was isolated from the thymus of euthanized mice with the RNeasy Mini Kit (Qiagen Canada, Toronto ON). RNA samples were sequenced using Truseq Stranded Total RNA Library Prep Kit and Illumina HiSeq4000 PE100 (Illumina, San Diego, CA) by the Genome Quebec Innovation Centre. The BAM files were trimmed using Trimmomatic [22]. Counts of annotated transcripts were performed using Salmon [23]. Differentially expressed genes were determined using DESeq2 [24]. RNA-sequencing results were visualized using Integrated Genomics Viewer [25]. Heat maps and enrichment plots were generated using the heatmap function in RStudio v1.3 or Gene Set Enrichment Analysis (GSEA) [26] using gene sets derived from the Panther database, or manually through literature search. Pathways were identified as potential targets of NAC using the Gene Ontology Resource [27], or the Database for Annotation, Visualization and Integrated Discovery (DAVID) and Kyoto Encyclopedia of Genes and Genomes (KEGG) [28].

### 2.5. PTEN mobility shift assay

The preleukemic cell line SepMBr was generated as described above. Whole-cell lysates were prepared from 5x10^6^ cells treated with *N*-acetylcysteine (MilliporeSigma A7250) for 4 hours followed by stimulation/oxidation with 1 mM hydrogen peroxide (Fisher Scientific Canada, Mississauga ON) for 5 minutes. Cells were washed and resuspended in PBS with 10% trichloroacetic acid (MillipreSigma T9159), sonicated, and washed again in 100% acetone. To preserve PTEN protein in its reduced state, protein precipitates were resolubilized in non-reducing lysis buffer (2% SDS, 50 mM Tris-HCl pH 6.8, 10% glycerol, 0.1% bromophenol blue) containing 40 mM *N*-ethylmaleimide (MilliporeSigma E3876). Samples were denatured at 100°C and separated on 10% SDS-PAGE gels with BLUeye Prestained Protein Ladder (FroggaBio, Concord ON). Protein was then semi-dry transferred to a PVDF membrane and incubated in Li-Cor Intercept^®^ Blocking Buffer (Cedarlane, Burlington ON). Immunoblotting was performed with monoclonal anti-PTEN rabbit IgG primary antibody (Cell Signalling Technology 9188S, Danvers MA) at 4°C overnight and goat anti-rabbit IgG secondary antibody (Li-Cor 926-32211) for 1 hour at room temperature. Blots were visualized with the Odyssey CLx Infrared imaging system at 800 nm (Li-Cor).

### 2.6. Sanger Sequencing

Genomic DNA was prepared from single cell suspensions of thymic leukemias using the Wizard Genomic DNA Purification Kit (Promega, Madison WI). PCR amplification of the *Jak3* pseudokinase domain was performed using MyTaq HS Red Master Mix (FroggaBio) and primers FWD: 5’- GCA TCA TGG TGC AGG AAT TTG T -3’ and REV: 5’- CGT TGA GGT CTC TGA GGA TAG C -3’. Sanger sequencing was performed using primers R653H FWD: 5’- AGG AGA GTT TGC TCA CAA C -3’ and R653H Seq REV: 5’- ACT TTC CAG GCT CAG CAC AG -3’.

### 2.7. Statistics

Statistical analysis was performed using Prism 8.4.3 (Graphpad, San Diego, CA). Individual statistical tests are named in the figure legends.

## 3. Results

### 3.1. Administration of 1.0 g/L N-Acetylcysteine does not confer a survival advantage

Mb1-CreΔPB mice develop leukemia at 100% incidence with a median time to euthanasia of 18 weeks [16,17]. Mb1-CreΔ1PB leukemias have high levels of ROS that results in cellular DNA damage [17]. We hypothesized that Mb1-CreΔPB mice provided with drinking water containing the antioxidant N-acetylcysteine (NAC) would develop B-ALL and display signs of illness later than Mb1-CreΔPB mice fed control water. For the first experiment, a dose of 1.0 g/L was used in order to alleviate potential toxicity [29]. A total of 30 mice were included in this experiment: 15 mice on control water and 15 on NAC water **(Supplementary Table 1)**. All mice consumed an average 6 mL of water per day, irrespective of the treatment group. The mice on control water and the mice on NAC water both developed leukemia and were euthanized upon first signs of illness including rapid, laboured breathing and hunched appearance, at a median age of 16.14 weeks **(Fig. 1A-C).**

**Fig. 1.**
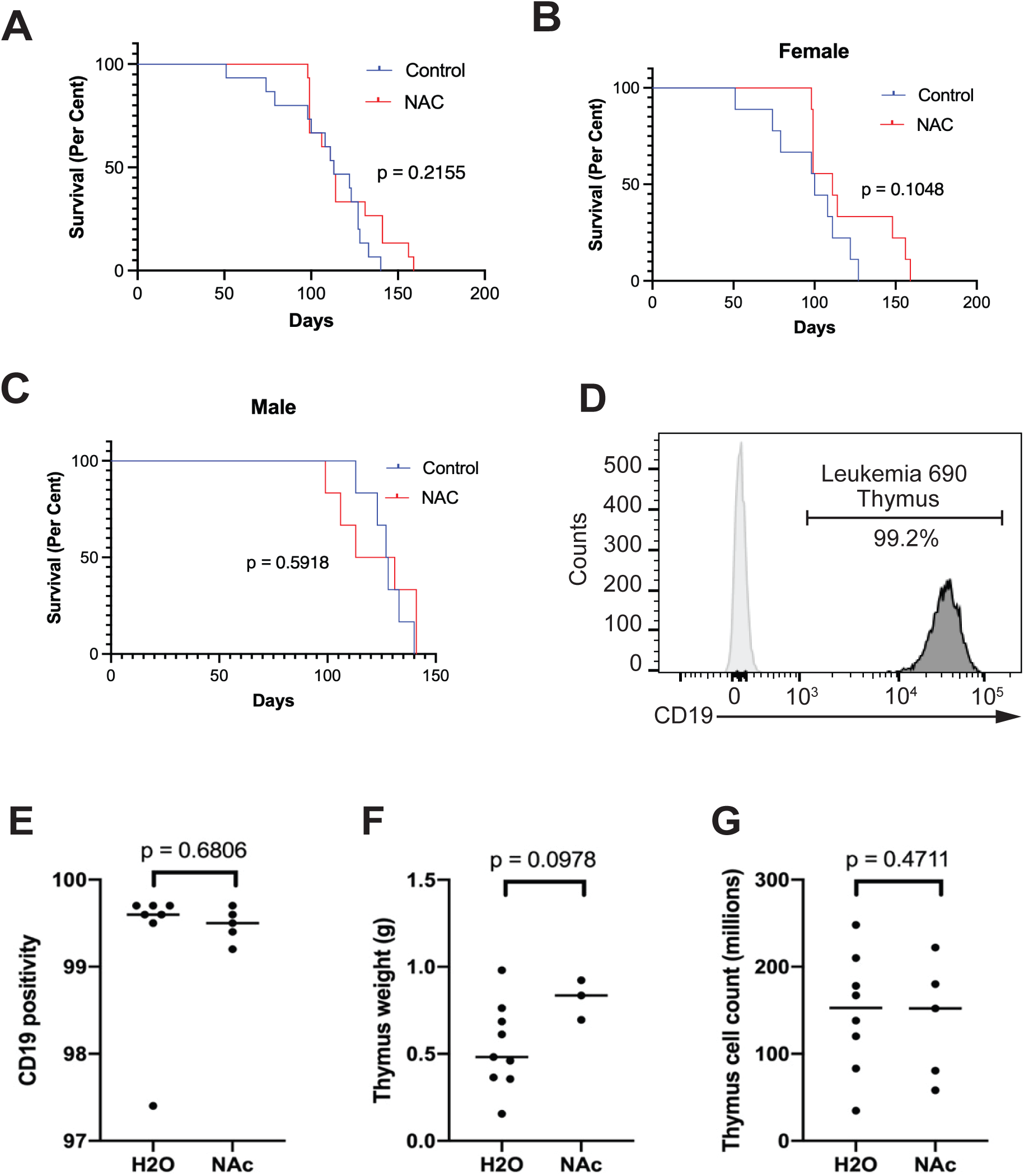
No increased survival in Mb1-CreΔPB mice administered 1 g/L N-acetylcysteine through drinking water. **(A)** Kaplan-Meier survival curve of Mb1-CreΔPB mice administered either 1 g/L N-acetylcysteine (NAC) in drinking water or control water starting at weaning (21 days of age). Log-rank (Mantel-Cox) test, n=15 per group. **(B, C)** Kaplan-Meier survival curve of mice from **(A)** separated by sex. Log-rank (Mantel-Cox) test, female n=9 per group, male n=6 per group. **(D)** Frequency of CD19^+^ B-ALL cells in thymus of mouse 690 from the water control group. **(E)** No difference in frequency of CD19^+^ thymocytes upon euthanasia (unpaired t test, control n=7, NAC n=5). **(F)** No difference in thymus weights upon euthanasia (unpaired t-test, control n=9, NAC n=3). **(G)** No difference in thymus cell counts upon euthanasia (unpaired t-test, control n=9, NAC n=5).

In the Mb1-CreΔ1PB mouse model, mice become ill with dyspnea caused by an enlarged thymus that is extensively infiltrated with pre-B ALL cells [16]. The thymus of moribund Mb1-CreΔ1PB mice typically contains greater than 97% CD19^+^ B-ALL cells and is therefore a nearly pure source of B-ALL cells for analysis (**Fig. 1D**). A subset of the 30 mice used in this experiment had their thymuses removed for measurement of weight, cell count, and cellular CD19 positivity. The frequency of CD19^+^ B cells in the thymus was consistently above 97% for both water and NAC groups at euthanasia **(Fig. 1E, 1F)**. No significant difference was observed between treatment groups in any of these categories **(Fig. 1E, F**, and **G**). In summary, 1.0 g/L of NAC in drinking water did not alter the endpoint of disease in the Mb1-CreΔ1PB mouse model.

#### Female mice weigh significantly less than male mice, resulting in a higher NAC dose per body weight and weight correlates with survival on NAC

Experimental mice were weighed every 1-3 days to monitor health and to ensure that they did not become dehydrated. As expected, male mice weighed significantly more than female mice over the course of the study (**Fig. 2A**, **2B**). The water intake by males and females was not significantly different with an average daily consumption of 6.0 g for males and 5.6 g for females **(Fig. 2C)**. This corresponds to female mice weighing 75% that of male mice and female mice ingesting 90% of the water of male mice. There was a significant difference in the weights of NAC-treated and control-treated male mice, with NAC-treated male mice weighing less over the course of the study (**Fig. 2D**). When the analysis was restricted to weights at day 100 and beyond, both male and female NAC-treated mice weighed less than control mice (**Fig. 2E**). By 100 days, there was reduced survival in every group, leaving 6 control male mice, and 5 mice in each of the other three treatment/sex combinations.

**Fig. 2.**
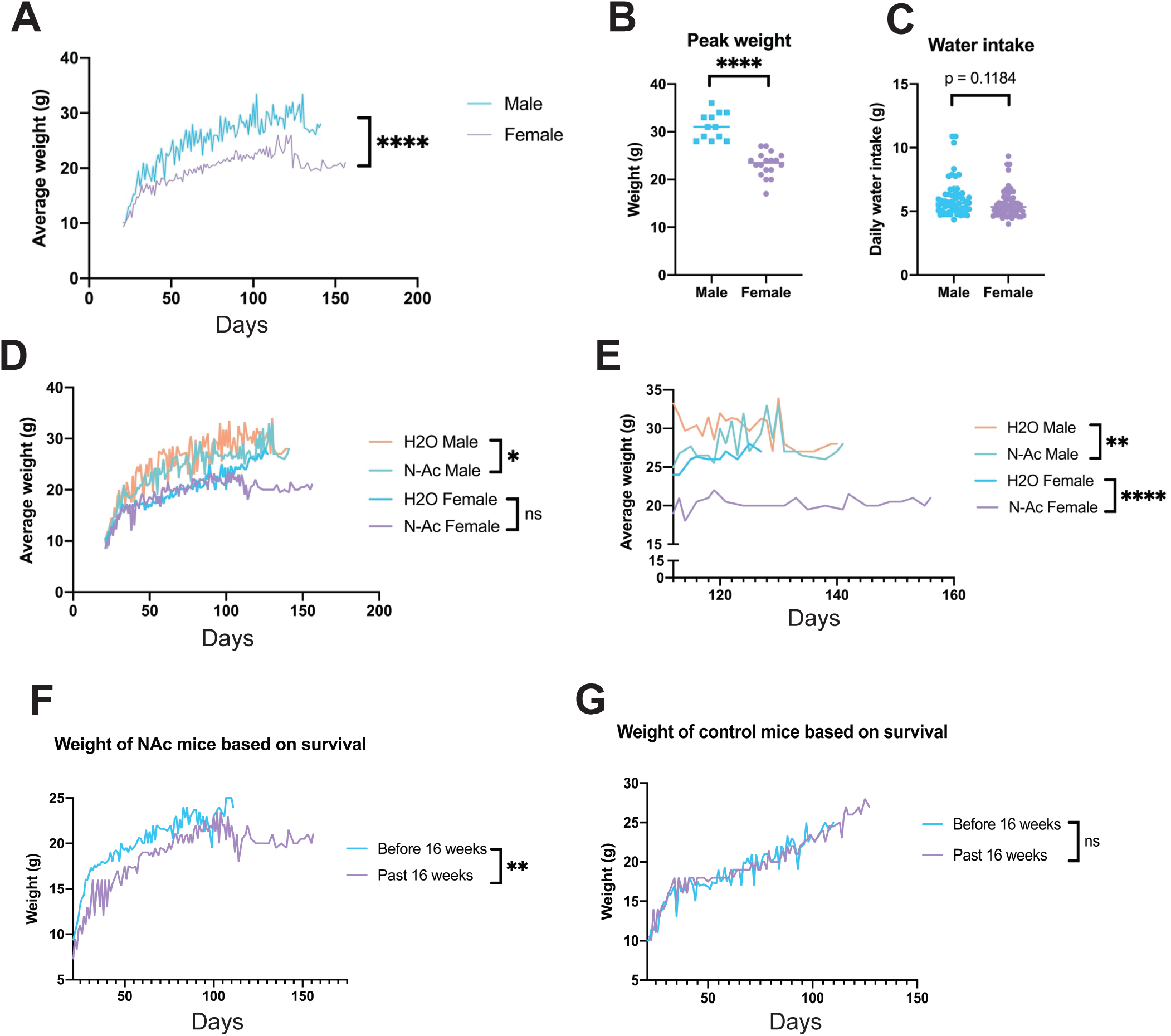
Increased survival in Mb1-CreΔPB mice with low body weights administered 1 g/L N-acetylcysteine. **(A)** Adult male mice weighed significantly more than female mice over the course of the study. **** p<0.0001 by unpaired t-test, male n=12, female n=18. **(B)** The peak weight reached by male mice is higher than the peak weight of female mice. **** p<0.0001, unpaired t-test, male n=12’ female n=18. **(C)** The amount of water ingested by male mice is not different than that ingested by female mice. Data points represent daily water intake averaged between 3 sets of cage mates between 21 and 77 days of age. Unpaired t-test, male n=6, female n=6. **(D)** Male mice receiving 1g/L N-acetylcysteine (NAC) through drinking water weigh less than male mice on regular drinking water. Unpaired t-test, control male n=6, NAC male n=6, control female n=5, NAC female n=5. **(E)** Male and female mice surviving more than 100 days receiving 1g/L NAC through drinking water weigh less than mice on control drinking water. ** p<0.01, **** p<0.0001 by unpaired t-test, control male n=6, NAC males n=5, control females n=5, NAC females n=5. **(F)** Female mice on NAC drinking water that survive past 16 weeks (100 days) weighed less than female mice on NAC drinking water that require euthanasia before 16 weeks. ** p<0.01 by unpaired t-test. **(G)** Female mice on control drinking water weighed the same whether they survived past 16 weeks or not.

When NAC-treated female mice were split into groups that survived less than or longer than 16 weeks (112 days), a significant difference was observed in their weights, with mice living longer than 16 weeks weighing significantly less over the entire course of treatment **(Fig. 2F)**. This difference was not observed for control female mice surviving less than or longer than 16 weeks **(Fig. 2G)**. This result suggested that differential survival was due to mice with low body weight receiving a higher effective dose of NAC.

### 3.2. Administration of 6.5g/L N-Acetylcysteine does not confer a survival advantage

Observed differences between male and female survival on 1.0 g/L NAC, as well as differences in survival between different weight classes of female mice suggested that the 1g/L NAC dose might be too low at adult body weight to cause differences in life span and suggested that a higher dose might be more effective in preventing B-ALL development. To circumvent toxicity when mice weighed less than 15 g [29], Mb1-CreΔPB mice were administered 1 g/L NAC water starting at day 21 after birth and then switched to 6.5 g/L when they reached a body weight of 15 g. It was observed that male mice reached 15 g in body weight at 5 weeks of age, whereas female mice reached 15 g by 6 weeks and thus these were the selected timepoints for dosage increase (**Fig. 3A**). Ten mice including both sexes were placed on NAC water and 10 mice of both sexes were maintained on regular drinking water (**Supplementary Table 2**). However, even at this increased NAC dosage, there were no significant differences in time to illness and euthanasia **(Fig. 3B)**. There were no significant differences in the frequency of CD19^+^ leukemia cells in the thymus, thymic weight, or thymic cellularity (**Supplementary Table 2**).

**Fig. 3.**
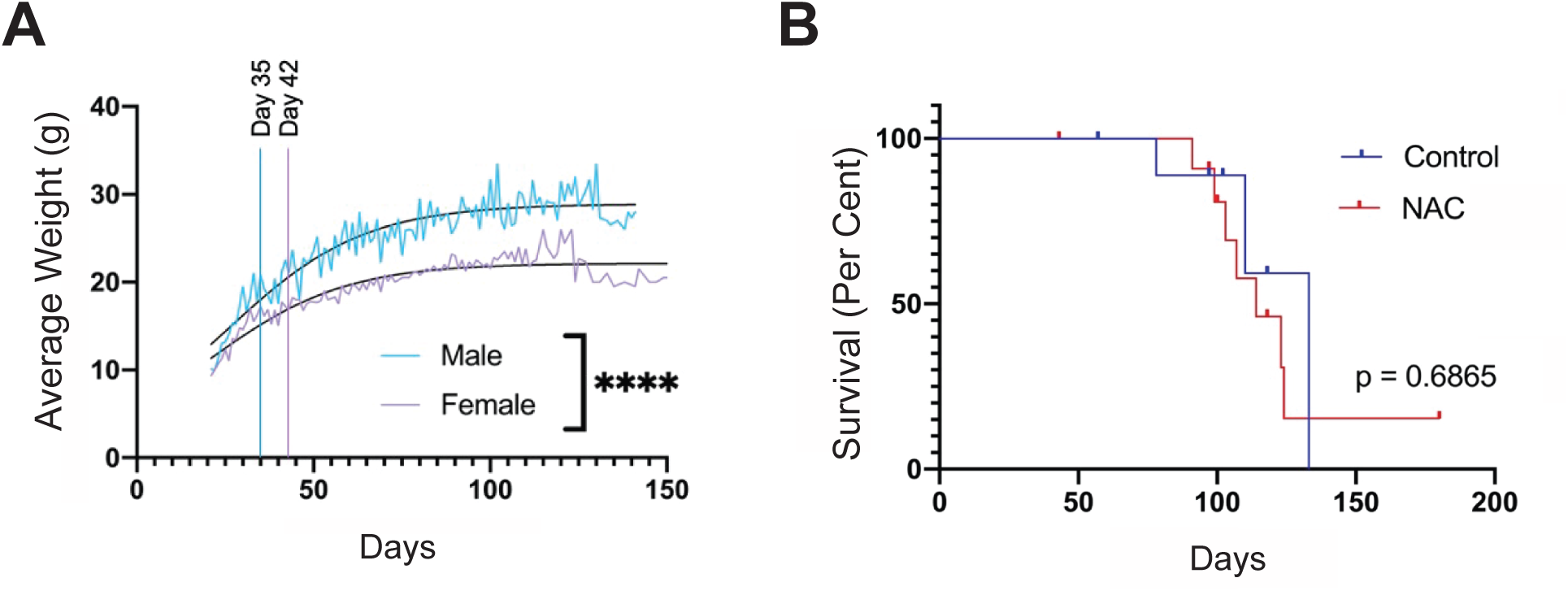
No increased survival in Mb1-CreΔPB mice administered 6.5 g/L N-acetylcysteine through drinking water. **(A)** Adult male mice weighed significantly more than female mice over the course of the study. **** p<0.0001 by unpaired t-test, male n=8, female n=20. Days 35 and Day 42 are indicated at which male and female mice reached 15 g of body weight. **(B)** Kaplan-Meier survival curve of Mb1-CreΔPB mice administered either 6.5 g/L N-acetylcysteine (NAC) in drinking water or control reverse-osmosis tap water starting at day 21 of age. Log-rank (Mantel-Cox) test, n=14 per group (10 females, 4 males).

### 3.3. Administration of 1.0 g/L of N-acetylcysteine results in delayed leukemia progression at 11 weeks of age

CD19^+^ leukemias are consistently detectable in the thymus of Mb1-CreΔ11PB mice by 11 weeks of age [16]. To determine if there were differences in the time of leukemia onset, the 11-week timepoint was selected to euthanize mice maintained on control or 1.0 g/L NAC drinking water. A total of 16 mice were included in this experiment **(Supplementary Table 3)**. At 11 weeks, mice maintained on 1.0 g/L NAC water had significantly lower frequencies of CD19^+^ leukemia cells within the thymus than control mice **(Fig. 4A)**. There was no significant difference observed between treatment groups in terms of thymus weight and thymus cell count **(Fig. 4B and C)**. In summary, 63% of control mice had infiltration of CD19^+^ leukemia cells into the thymus at 11 weeks, compared to 13% of NAC-treated mice, suggesting that NAC treatment resulted in delayed leukemia progression at this time point.

**Fig. 4.**
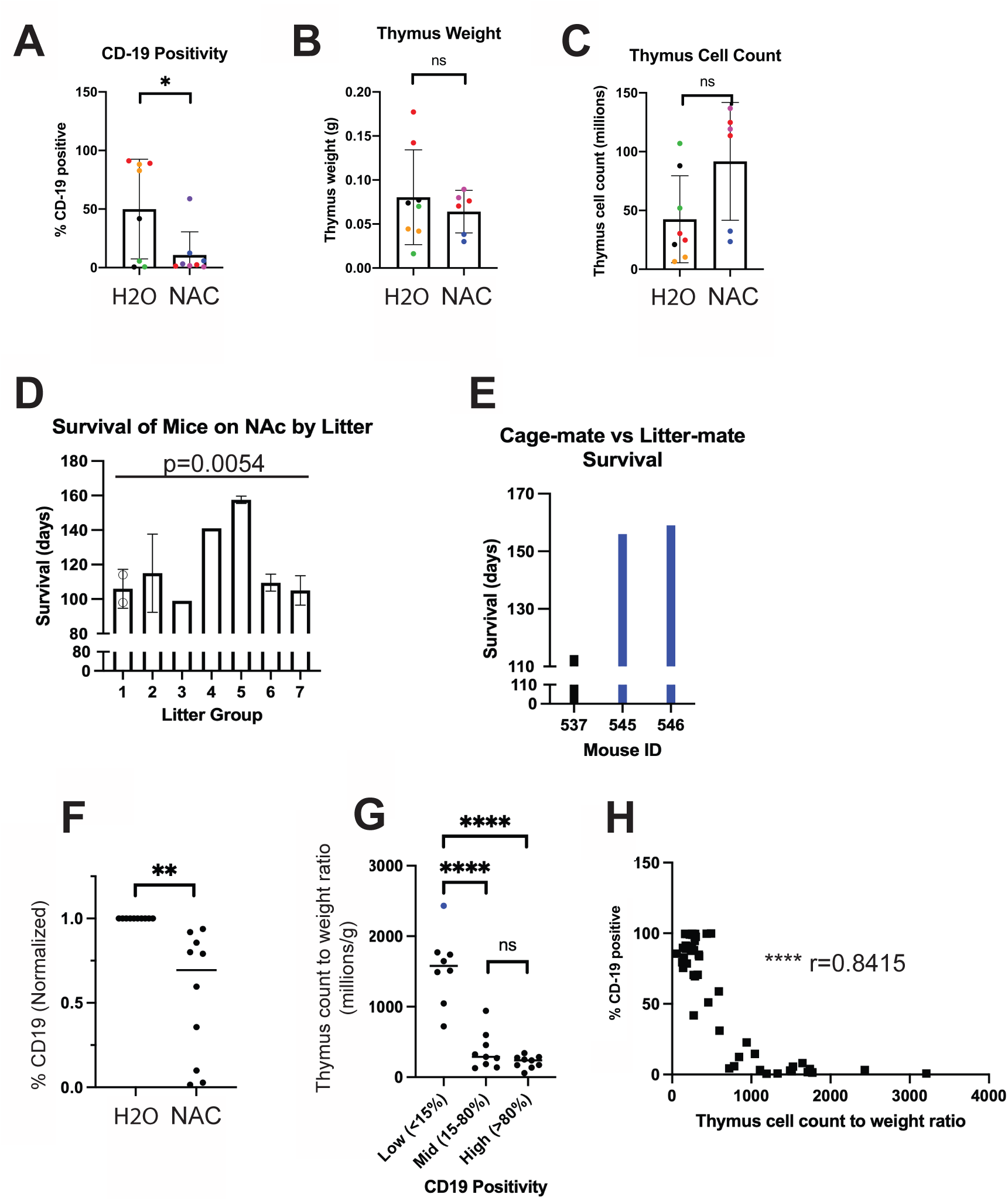
Reduced leukemia progression at 11 weeks of age in Mb1-CreΔPB mice administered 1 g/L N-acetylcysteine. **(A)** Mice administered 1g/L N-acetylcysteine (NAC) had lower CD19+ B cell frequencies in the thymus at 11 weeks compared to mice administered control drinking water. * p<0.05 by unpaired t-test, control n=8, NAC n=8. **(B)** No difference between thymus weight at 11 weeks in mice administered 1 g/L NAC compared to control drinking water. Unpaired t-test, control n=8, NAC n=8. **(C)** No difference between thymus cell counts at 11 weeks in mice administered 1 g/L NAC compared to control drinking water. Unpaired t-test, control n=8, NAC n=8. **(D)** Mb1-CreΔ1PB mice from the same litter tend to develop leukemia at similar timepoints. Of 7 litter groups with two mice each, four had two mice develop illness within 7 days of each other, and 5 within two weeks. P=0.0054 by one-way ANOVA. **(E)** Representative littermate data. One cage group from (A) contained two female mice from the same litter (545 and 546) and one female mouse born the same day from a different litter (537). Mice 545 and 546 developed illness at days 156 and 159, while mouse 537 developed illness at day 114. **(F)** Littermate pairs of mice administered 1g/L NAC water had lower CD19^+^ B cell frequencies in the thymus at 11 weeks of age than control mice. ** p<0.05 by one sample t and Wilcoxon test, n=10 in each group. **(G)** Ratio of thymus counts to thymus weight binned by CD19 positivity. **** p<0.0001 by one-way ANOVA. **(H)** Correlation of CD19 positivity (y-axis) with thymus cell count to weight ratio (x-axis). **** p<0.0001 by Pearson Correlation Coefficient.

In the initial 1.0 g/L experiment, mice were placed in either treatment or control groups with 2-3 littermates in each cage. There was a tendency for mice from the same litter to develop leukemia at similar timepoints compared to mice from different litters but in the same treatment group (**Fig. 4D**). Of the 7 littermate groups with two mice each, 4 had the two mice develop illness within 7 days of each other, and 5 within two weeks of each other. One particular cage of mice housed three mice treated with NAC: two mice from one litter (mouse tag 545 and 546) and one mouse from a different litter (tag 537) but born on the same day. All three mice were housed together for the duration of the experiment. The two mice from the same litter developed illness within three days of each other, while the individual mouse from the separate litter developed illness 6 weeks before the earlier of the two litter-mates **(Fig. 4E)**. These results suggest a genetic or epigenetic influence on the progression of leukemia in the Mb1-CreΔPB mouse model.

To control for the similar timing of leukemic progression within littermate groups, a second 11-week experiment was conducted with 1.0 g/L NAC drinking water and litter-matched mice. When litters had three or more Mb1-CreΔPB mice of the same sex, two mice were randomly selected for NAC treatment and the remaining 1 or 2 mice were placed into the control water group **(Supplementary Table 4)**. The mice were euthanized at 11 weeks of age and their thymuses were analyzed for CD19 positivity. NAC-treated mice were analyzed relative to their control littermates. The NAC-treated mice had significantly lower CD19 positivity than their control littermates, and no NAC-treated mouse group had a higher CD19 positivity than their control littermates **(Fig. 4F)**. Finally, the thymus cell count to weight ratio was analyzed against CD19 positivity for all mice in both 11-week experiments. A clear trend showing increased CD19 positivity and lower thymus cell count to weight ratios was observed **(Fig, 4G, 4H)**. In summary, these results showed that NAC treatment significantly delayed leukemia progression at the 11-week timepoint when controlled for epigenetic or genetic effects within littermate groups.

### 3.4. Leukemias from mice on NAC and control water have significant differences in gene expression

To determine whether NAC treatment led to differences in gene expression, a subset of mice from the 1.0 g/L trial had RNA prepared from their thymuses upon euthanasia, containing > 97% of CD19^+^ pre-B-ALL cells (**Fig. 1E**). RNA-sequencing was performed on 8 water control and 8 NAC-treated leukemias using paired-end Illumina sequencing. Count tables for annotated mRNA transcripts were determined using Salmon [23]. One leukemia (597) was excluded from further analysis because gene expression suggested that the leukemia was T-ALL rather than B-ALL (data not shown). Principal component analysis performed on Log_2_-transformed count data suggested that gene expression from leukemias in NAC-treated mice were more similar to one another than leukemias from control water-treated mice (**Fig. 5A**).

**Fig. 5.**
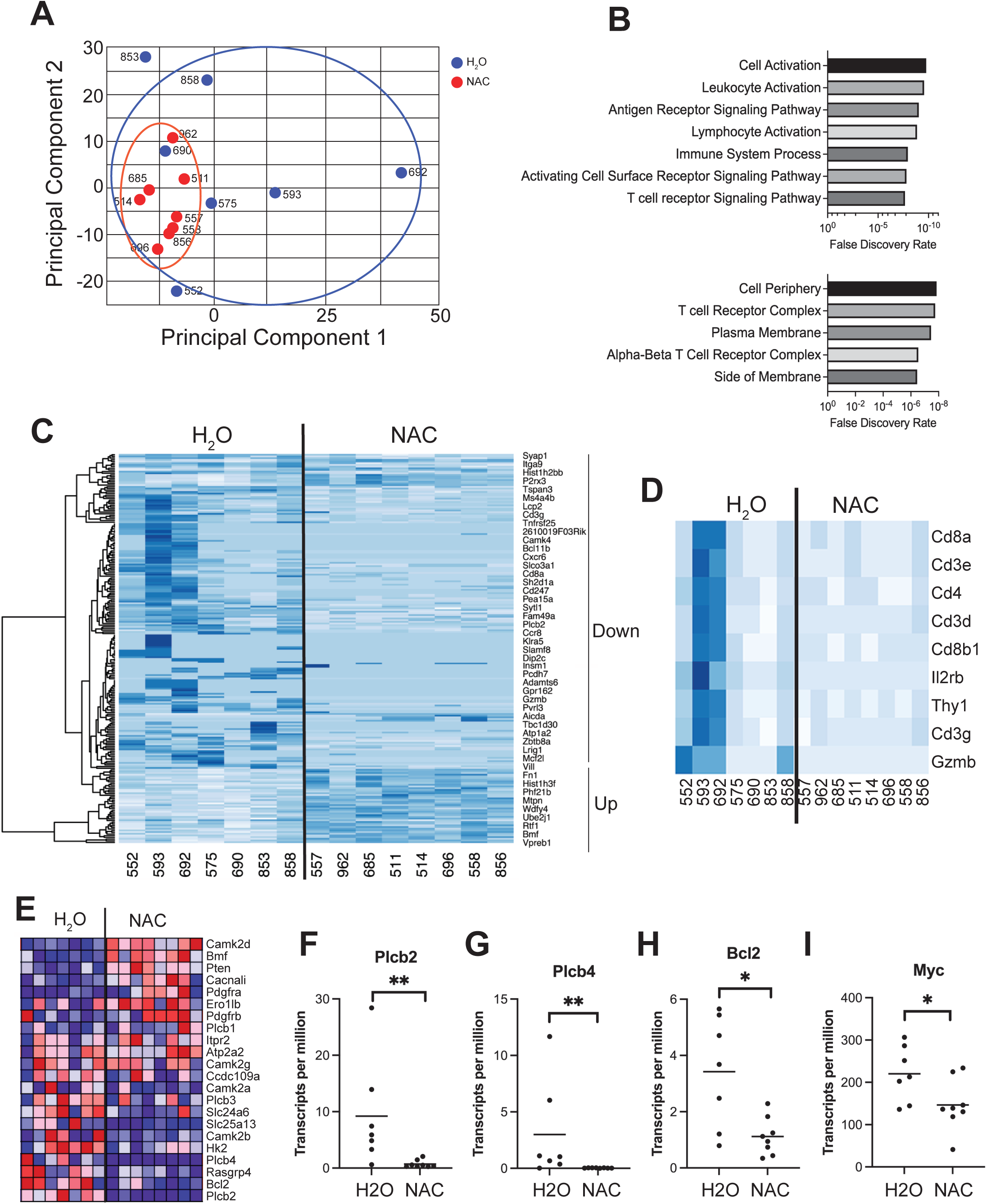
Differences in gene expression between leukemias from mice treated with 1g/L N-acetylcysteine and leukemias from control mice. **(A)** Principal component analysis of RNA-sequencing count data from leukemias from mice treated with N-acetylcysteine (NAC n=8, red) or control drinking water (blue, n=7). Circles indicate grouping of samples. Mouse identification numbers are indicated. **(B)** Gene Ontology analysis of genes differentially expressed between leukemias from mice treated with NAC or control drinking water. Top panel, top 7 biological pathways ranked by low false discovery rate. Bottom panel, top 5 cellular components ranked by low false discovery rate. **(C)** Heatmap analysis of gene expression differences. Unsupervised clustering of RNA-sequencing count data for 254 genes was performed. Hierachical clustering is shown on left, representative gene names and up- or down-regulation is shown on right. **(D)** Heatmap analysis of T cell specific transcripts indicated on right. **(E)** Heatmap generated from Gene Set Enrichment Analysis for the calcium signaling gene set. **(F-I)** Count data for four representative genes downregulated by NAC (*Plcb2*, *Plcb4*, *Bcl2*, and *Myc*) expressed as transcripts per million base pairs. * p<0.05, ** p<0.01 by unpaired t-test.

Using DESeq2 [24] 254 genes were identified as being differentially expressed between treatment groups (p<0.05, **Supplementary Dataset 1**). We observed that 61/254 genes were upregulated while 193/254 genes were downregulated in the NAC group relative to the control water group. Gene Ontology analysis showed that altered biological pathways included *Cell Activation*, *Leukocyte Activation*, *Antigen Receptor Signaling*, and *Lymphocyte Activation* (**Fig. 5B, top panel**). Altered Cellular Components included *Cell Periphery*, *T cell Receptor Complex*, *Plasma Membrane*, and *Alpha-Beta T Cell Receptor Complex* (**Fig. 5B, lower panel**). Heatmap analysis of Log_2_-transformed count data showed substantial gene expression differences between groups, with both up-regulated and down-regulated transcripts (**Fig. 5C, Supplementary Dataset 2**). Examination of T cell-specific transcripts showed that leukemias from control mice had higher levels than NAC-treated mice, with most of the T cell-specific transcripts being expressed in leukemias 593 and 692 (**Fig. 5D**). This result suggested either a difference in immune response to the tumour, or a difference in B-ALL cell frequency in the thymus in these samples.

The *calcium signaling* pathway was identified as a significantly differentially expressed biological pathway using DAVID. The most downregulated genes identified by DESeq2 were *Plcb2* and *Plcb4* (encoding phospholipase C beta 2 and 4), which are implicated in calcium signaling. Using Gene Set Enrichment Analysis (GSEA) and a gene set involved in calcium signaling between the endoplasmic reticulum and mitochondria that regulates apoptosis in high ROS concentrations, there was clear dissimilarity between the treatment groups **(Fig. 5E)**, with genes including *Plcb2*, *Plcb4*, *Bcl2*, and *Myc* downregulated in leukemias from NAC-treated mice **(Fig. 5F-I)**. In summary, leukemias from NAC-treated mice had a substantially different pattern of gene expression than leukemias from control mice, suggesting altered clonal evolution.

### 3.5. Increased Jak3 R653H mutation frequency in leukemias from NAC-treated mice

We previously reported that activating mutations in *Jak1* and *Jak3* encoding Janus Kinases 1 and 3 are frequent in the Mb1-CreΔ1PB mouse leukemia model [16,17]. Integrated Genome Viewer (IGV) was used to examine RNA sequencing data for *Jak1* and *Jak3* mutations present in nucleotides with >500 reads and in which the single nucleotide variant was present in 10% or more of the reads. An example of *Jak3* R653H mutation identification based on RNA-sequencing is shown in **Fig. 6A**. A summary of the mutations observed can be found in **Fig. 6B**. All NAC-treated leukemias had a mutation in either *Jak1* or *Jak3* (**Fig. 6B**). Five different mutations were observed in *Jak1*, 4 of which were located in the pseudokinase domain, and the remaining being found between the SH2 and kinase region (**Fig. 6B, C**). NAC treated mice had higher frequencies of *Jak3* R653H mutation (6/8) than control water treated mice (2/7), and in those leukemias that did harbour the mutation, NAC mice had a higher *Jak3* R653H variant allele frequency (**Fig. 6D**). All R653H mutations were G->A mutations in the codon C<u>G</u>T (R) -> C<u>A</u>T (H). Next, PCR amplification followed by Sanger sequencing was used to confirm and extend the RNA-sequencing results. Interestingly, *Jak3* R653H mutation frequency was reduced in thymic CD19^+^ cells at 11 weeks after treatment with 1.0 g/L NAC drinking water (**Fig. 6E**) but increased in leukemias from mice euthanized due to illness at a median 16 weeks after treatment (**Fig. 6F**). An example of a sequence chromatogram for *Jak3* R653H mutation from leukemia 856 is shown in **Fig. 6G**. In summary, NAC treatment of Mb1-CreΔ11PB mice results in increased *Jak3* R653H mutation frequency in the leukemias that develop by median 16 weeks of age.

**Fig. 6.**
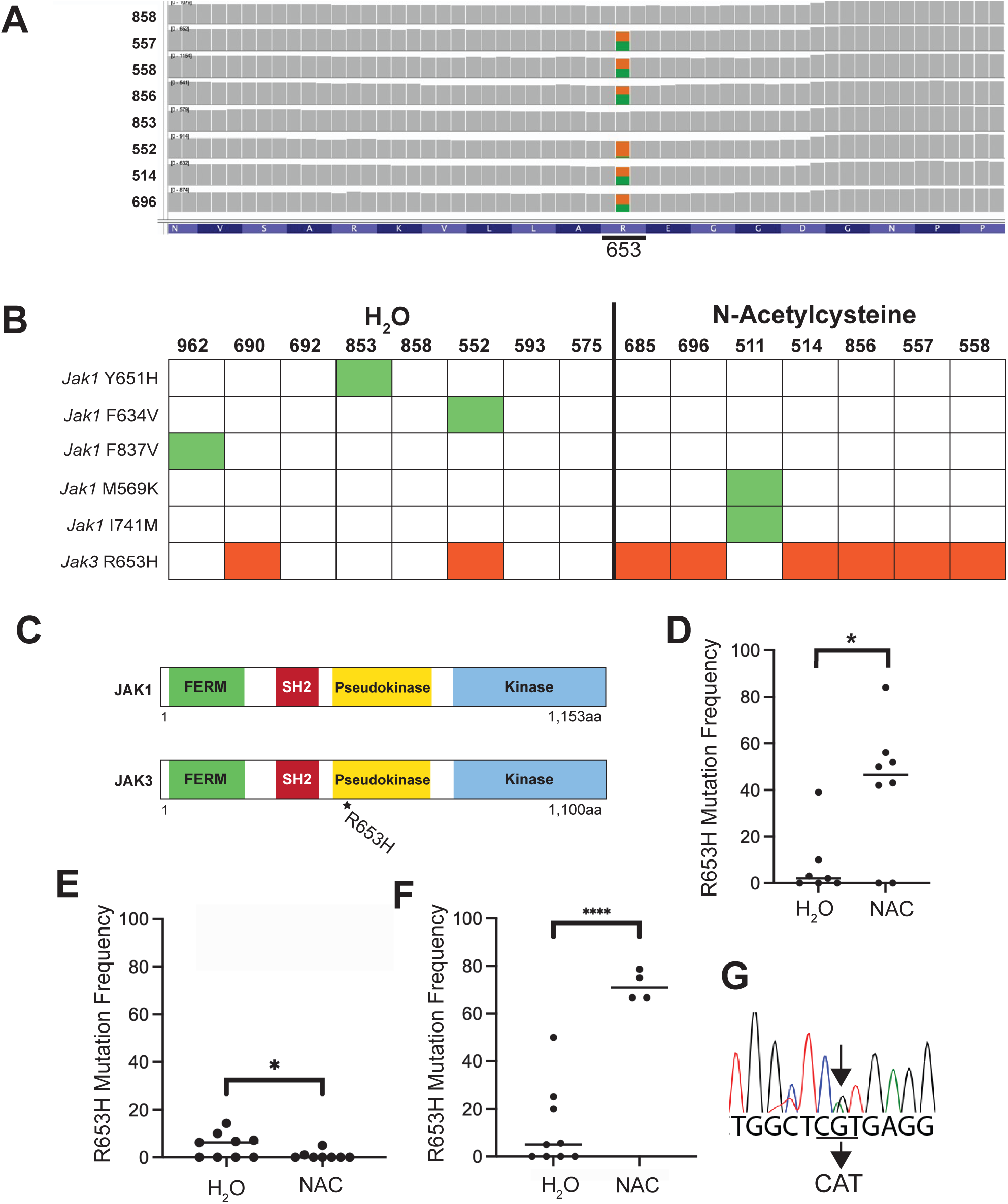
*Jak3* R653H mutation frequency is increased in mice treated with N-acetylcysteine. **(A)** Representative examination of RNA-sequencing data using integrated genome viewer (IGV). Mouse numbers are indicated on left. Amino acid position of R653 is indicated on bottom. **(B)** Heat map to indicate frequencies of *Jak1* and *Jak3* mutations. *Jak1* mutations are indicated on the left and in green. *Jak3* mutations are indicated in red. **(C)** Schematic of JAK1 and JAK3 protein structures to show functional domains. Location of R653H is indicated. **(D)** Frequency of *Jak3* R653H mutation is increased in leukemias from Mb1-CreΔPB mice placed on NAC drinking water compared to mice on control drinking water. Frequency was calculated from RNA-sequencing data as shown in (a). **(E)** Frequency of *Jak3* R653H mutation is decreased at 11 weeks in leukemias from Mb1-CreΔPB mice placed on NAC drinking water compared to mice on control drinking water. Mutation frequency is calculated from Sanger sequencing. **(F)** Frequency of *Jak3* R653H mutation is increased at euthanasia weeks in leukemias from Mb1-CreΔPB mice placed on NAC drinking water compared to mice on control drinking water. Mutation frequency is calculated from Sanger sequencing. **(G)** Representative Sanger sequencing chromatogram from leukemia 856. G->A mutation is indicated.

### 3.6. Redox regulation of PTEN by H_2_O_2_ and N-acetylcysteine

Genes upregulated in NAC-treated leukemias compared to control water leukemias included *Vpreb1*, *Bmf*, and *Pten* (**Fig. 7A-C**). *Vpreb1* encodes a component of the pre-B cell receptor surrogate light chain protein and is highly expressed in pre-B-ALL cells in the Mb1-Cre-Δ11PB mouse model [30]. *Bmf* encodes Bcl-2-modifying factor and functions as an apoptotic activator [31]. This result, along with decreased transcript levels of *Bcl2* encoding the anti-apoptotic protein Bcl-2 (**Fig. 5G**) suggested altered apoptosis in NAC-treated leukemias.

**Fig. 7.**
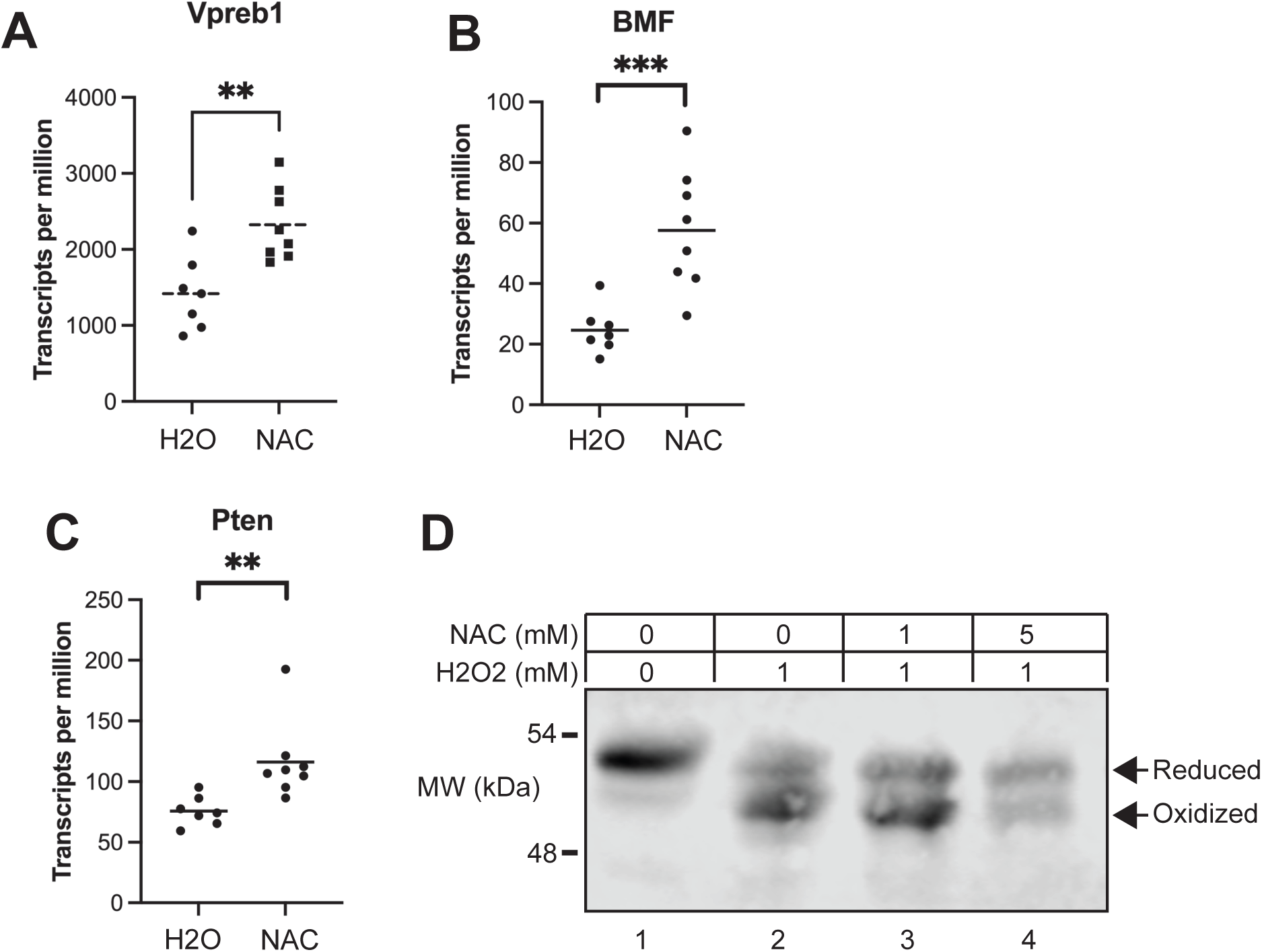
Reduction of PTEN by N-acetylcysteine. **(A-C)** Count data for three representative genes upregulated by NAC (*Vpreb1*, *Bmf*, *Pten*) expressed as transcripts per million base pairs. * p<0.05, ** p<0.01 by unpaired t-test. **(D)** PTEN protein is oxidized by ROS and reduced by NAC. Lysates were prepared from 973 leukemia cells treated with 1 mM H2O2 for 4hr, or with 1 mM H2O2 and 1 or 5 mM NAC for 4 hr before separation on nonreducing SDS-PAGE gel. Immunoblot was performed using anti-PTEN antibody. Reduced and oxidized forms of PTEN are indicated.

*Phosphatase and tensin homolog deleted on chromosome ten* (PTEN) functions as a lipid phosphatase on the signaling mediator PtdIns(3,4,5) to regulate cell proliferation and metabolism [32]. Interestingly, although PTEN is considered a tumor suppressor that is frequently mutated in cancer, *Pten* is rarely mutated in pre-B-ALL [33]. Pre-B cells are uniquely sensitive to induction of apoptosis by PtdIns(3,4,5)-induced signaling, and PTEN is essential to protect pre-B cells from cell death induced by this pathway [33]. PTEN activity is modulated by cellular Redox status as oxidation of cysteine resides at its active site reversibly inactivates its activity [34]. We therefore hypothesized that PTEN activity might be regulated by the high levels of ROS in Mb1-CreΔPB leukemias [17]. We observed that NAC-treated leukemias expressed higher levels of *Pten* mRNA transcripts compared to control leukemias, suggesting up-regulation of *Pten* as a compensatory effect in response to NAC (**Fig. 7C**).

To determine if PTEN active-site oxidation can be regulated by NAC in mouse pre-B cells, we used an assay in which PTEN mobility in a nonreducing SDS-PAGE gel is determined by immunoblot following oxidation or reduction [35]. The IL-7-dependent preleukemic pre-B cell line SeptMBr isolated from Mb1-CreΔPB mice was treated with H_2_O_2_ to induce oxidization of PTEN and treated with NAC to determine if this could reduce PTEN. As shown in **Fig. 6D**, treatment of pre-B cells with H_2_O_2_ oxidized PTEN to the more rapidly migrating form as detected by this assay. Treatment of pre-B cells with 1 or 5 mM of NAC reversed the oxidation of PTEN by 1 mM H_2_O_2_. This result suggests that NAC opposes the effect of ROS on PTEN oxidation state, and therefore likely alters PTEN activity. NAC would therefore be expected to maintain PTEN in its reduced, active state to promote pre-B-ALL survival and proliferation [32].

## 4. Discussion

In this study, we experimentally tested the agent N-acetylcysteine (NAC) for its ability to alter leukemia progression in the Mb1-CreΔ1PB mouse model. Mice treated with 1.0 g/L of NAC in drinking water were found to have reduced frequencies of CD19^+^ B-ALL cells infiltrating the thymus at 11 weeks of age. However, mice treated with 1.0 g/L or 6.5 g/L NAC in drinking water had similar times to euthanasia as control mice by median 16 weeks of age. RNA-sequencing analysis showed that leukemias in mice treated with 1.0 g/L NAC had an altered pattern of gene expression compared to control mice. Leukemias from NAC-treated mice had 100% incidence of activating pseudokinase domain mutations in *Jak1* or *Jak3*. Activating R653H *Jak3* mutations were increased in frequency in NAC-treated mice compared to control mice. NAC opposed the effects of hydrogen peroxide on oxidation of PTEN to an inactive form, suggesting that redox status of PTEN played a role in disease progression. These results suggest that clonal evolution of leukemia in the Mb1-CreΔ1PB mouse model is altered by NAC-mediated modulation of reactive oxygen species, ultimately selecting for more aggressive tumours.

In the initial 1.0 g/L trial, mice receiving NAC were observed to have reduced frequencies of CD19^+^ B-ALL cells in the thymus at 11 weeks of age. Eleven weeks was chosen as an intermediate timepoint for leukemia progression based on previous work that showed that leukemia cells were consistently detectable in the thymus of Mb1-CreΔPB mice by 11 weeks of age [17]. At 11 weeks, most littermate pairs or groups had similar CD19 levels, usually either low (<15%) or high (>80%) CD19 positivity. When CD19 positivity data was analyzed to control for littermates, NAC-treated mice had significantly less B-cell infiltration into the thymus at 11 weeks of age, indicating less tumour growth, regardless of sex.

Although NAC-treated mice had reduced CD19 positivity at 11 weeks after receiving 1.0 g/L doses, there was ultimately no survival difference between treated and control mice at 1.0 g/L or 6.5 g/L. This result suggests that the leukemias observed in the NAC-treated mice had delayed progression, but progressed more rapidly after 11 weeks of age. This suggests that treatment with NAC causes faster onset of signs of illness from the start of B cell infiltration into the thymus.

While there were no differences in the survival of mice on NAC compared to control water, there was a clear trend where mice weighing the least over their entire lifespan tended to survive the longest when consuming NAC water (**Fig. 2F**). This result suggested that a higher NAC dose promoted survival. However, the time to euthanasia was not extended when the NAC dose was increased to 6.5g/L. The increased dose may have either promoted tumour progression, as discussed below, or increased toxicity that skewed the results from what was predicted. A future dosing experiment testing multiple intermediate doses with smaller cohorts of mice may elucidate an optimal dosage for future experiments.

RNA-sequencing analysis showed that gene expression was significantly altered in leukemias from NAC-treated mice compared to control mice. Principal component analysis suggested that leukemias from NAC-treated mice were more uniform compared to leukemias from control mice. We speculate that because leukemia progression in NAC-treated mice was delayed, but later caught up to control mice, that these leukemias were more aggressive and originated from fewer preleukemic clones. Several biological pathways were different between NAC-treated leukemias and control leukemias at the gene expression level. Genes in the calcium-apoptosis signaling pathway were determined to be differentially expressed between NAC treatment and control groups. *Plcb2* and *Plcb4* were the two most differentially expressed genes reported by DESeq2 and were downregulated in NAC-treated leukemias compared to control leukemias. In contrast, genes like *Bcl2* which inhibit the action of IP3R [36] were also downregulated in NAC-treated mice. Genes that function to activate calcium movement into the mitochondria, signaling apoptosis - such as *Camk2d*, *Bmf*, *Pten*, *Pdgfrb*, *Plcb1-4*, *Itpr3*, and *Rasgrp4* [37] were upregulated in NAC-treated mice. Furthermore, some of the genes involved in inhibiting this pro-apoptotic pathway, such as *Slc24a6*, *Hk2*, and *Bcl2* [37], were downregulated in NAC-treated mice. These results suggest that tumour cells in NAC-treated mice have increased activation of pro-apoptotic signaling through the endoplasmic reticulum-mitochondria calcium release axis.

Janus Kinase-1 and 3 mutations are recurrent in the Mb1-CreΔ1PB mouse model of leukemia [16,17]. JAK1 Y651H, F634V, F837V, I741M, and JAK3 R653H mutations are all located in the Janus Kinase pseudokinase domain and are thus predicted to act as activating mutations [38]. The recurrent *Jak3* R653H mutation was shown to increase proliferation of leukemia cells in response to low concentrations of IL-7 [16]. This mutation is equivalent to the *Jak3* R657Q mutation frequently observed in human ALL [39]. *Jak3* R653H mutation frequency was increased in mice treated with NAC drinking water (**Fig. 6D-G**). 100% of NAC-treated mice had a mutation in the pseudokinase domain of either *Jak1* or *Jak3*, whereas not every control mouse had detectable pseudokinase JAK mutations. This result suggests NAC treatment places selective pressure on leukemia clonal evolution in the Mb1-CreΔ1PB mouse model such that leukemias with Janus Kinase mutations are favoured.

PTEN is widely considered to be a tumour suppressor, and is frequently mutated in human cancers [32]. However, survival of pre-B cells is uniquely dependent on PTEN activity relative to other body cell types [40]. As a consequence, *PTEN* is rarely mutated in pre-B-ALL [40]. PTEN functions as a lipid phosphatase on the signaling mediator PtdIns(3,4,5), high levels of which are toxic for pre-B cells [32,40]. Our results showed that *Pten* mRNA transcript levels were increased in leukemias from NAC-treated mice, and NAC could directly oppose the reversible inactivation of PTEN by oxidation of cysteine residues at its active sites (**Fig. 7**). Previous studies showed that leukemias in the Mb1-CreΔPB mouse model have high levels of ROS; and furthermore, these leukemias appear to be addicted to high levels of ROS since most antioxidant genes were downregulated, and H_2_O_2_ was mitogenic for cultured leukemia cells [17]. We speculate that because of high ROS levels, PTEN would be primarily present in an oxidized, inactive state in Mb1-CreΔPB leukemia cells [34,41]. Decreased PTEN activity would increase PtdIns(3,4,5)- induced cell death in leukemia cells, resulting in increased apoptosis. Reduction of PTEN by NAC would increase activity, resulting in reduced apoptosis. The observation of reduced *Bcl2*, increased *Bmf*, and increased *Pten* mRNA transcript levels (**Figs. 5H**, **7B**, and **7C**) are compatible with this idea.

In conclusion, we speculate on the following model (**Fig. 8**). Mb1-CreΔPB mice develop recurrent Janus Kinase driver mutations early in clonal evolution, resulting in increased JAK/STAT signaling and high levels of ROS. High ROS is mitogenic for leukemias by activating JAK/STAT signaling [14]. However, high ROS levels result in PTEN inactivation and increased apoptosis [41]. High levels of ROS also induce cellular DNA damage, leading to increased apoptosis, but potentially also to secondary driver mutations and increased clonal diversity (**Fig. 5A**, **Fig. 8**). Treatment of Mb1-CreΔPB mice with NAC might oppose JAK/STAT signaling and activate PTEN, leading to reduced proliferation, as evidenced by reduced leukemia at 11 weeks of age, and reduced apoptosis. However, NAC might counter-productively select leukemia clones with *Jak3* R653H driver mutations that would lead to higher levels of ROS, overcoming the antioxidant effect of NAC. Ultimately this would lead to the selection of leukemias with reduced clonal diversity, but increased aggressiveness (**Fig. 8**). This model is consistent with all observations made in this study. Ultimately this should lead to caution in use of N-acetylcysteine or other antioxidants in the treatment of leukemia, due to the highly complex interplay between beneficial and harmful effects of ROS on various cellular processes. Indeed, the results of antioxidant clinical trials for human cancers have been mixed and have not yet led to clear treatment strategies [42,43].

**Fig. 8.**
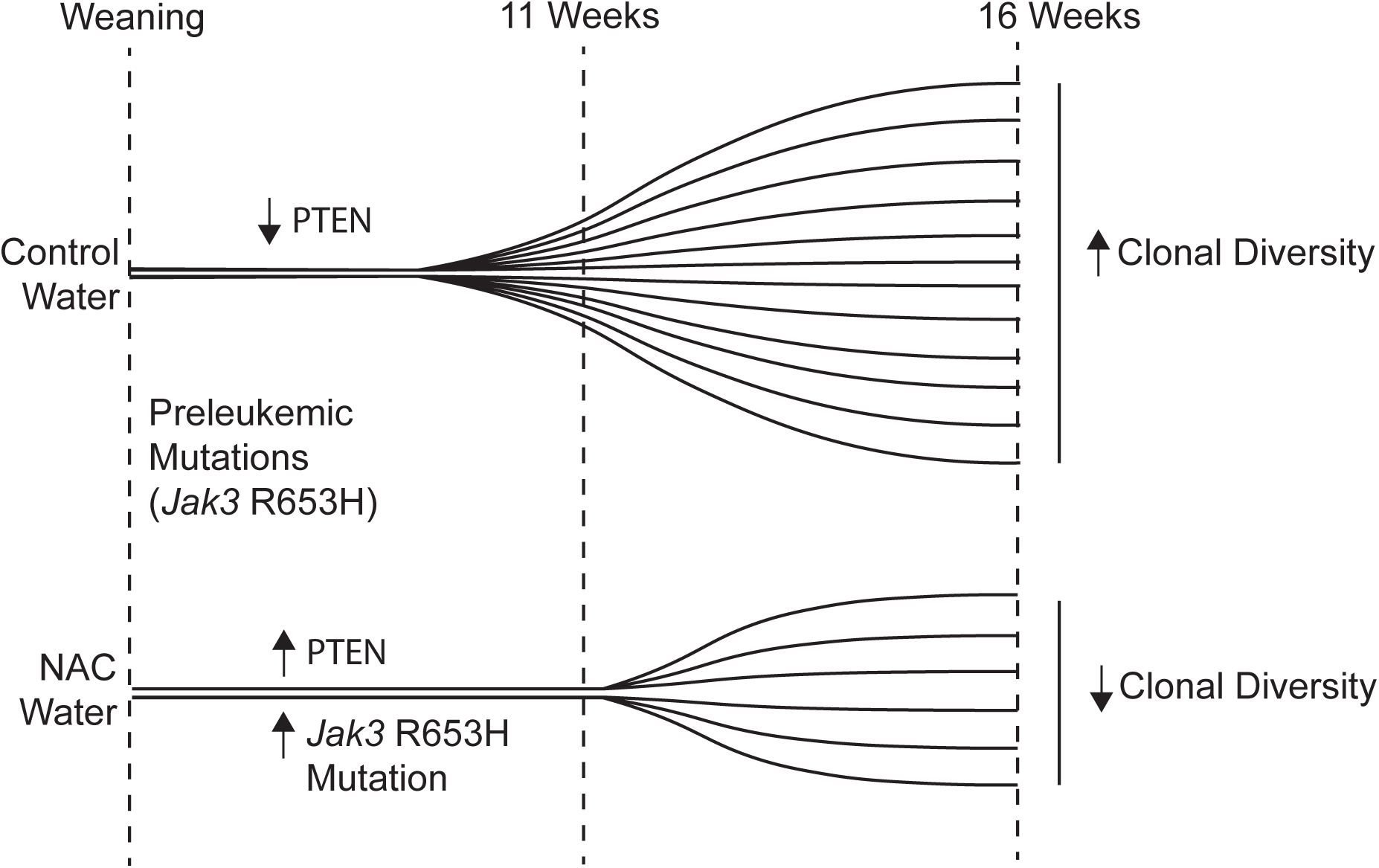
Model of disease progression of Mb1-CreΔPB leukemias in mice treated with control drinking water (top) or N-acetylcysteine drinking water (bottom). Disease is detectable at 11 weeks in control mice but not in NAC mice. Clonal diversity is increased in control leukemias but reduced in NAC mice.

## Supporting information

DESeq2 differentially expressed genes

Raw Salmon Counts

## Author Contributions

MPS was responsible for conducting literature searches, designing experimental protocols, performing experiments, and writing the manuscript. JI, MA, and BRO were responsible for designing experimental protocols and performing experiments. HR and MPS were responsible for generating the SeptMBr cell line. LSX was responsible for mouse care and colony management. RPD was responsible for conducting literature searches, designing experimental protocols, and writing the manuscript.

## Availability of Data and Materials

RNA Sequencing data is available from the Sequence Read Archive under BioProject Accession PRJNA988517. Reviewer link: https://dataview.ncbi.nlm.nih.gov/object/PRJNA988517?reviewer=v618hf9i922mf79ij32fa0j7q2

## Acknowledgements

We thank Kristin Chadwick from the London Regional Flow Cytometry core facility for assistance with Flow Cytometry. We thank Jenn Biltcliffe from the London Regional Genomics Centre for assistance with Sanger DNA sequencing. We thank Joshua Yi and Allanna Mackenzie for critically reading the manuscript. Funding: this work was funded by operating grants 142258 and 168995 from the Canadian Institutes of Health Research.

