## Supplementary Tables 1-4 for "N-Acetylcysteine alters disease progression and increases Janus Kinase mutation frequency in a mouse model of precursor B cell acute lymphoblastic leukemia"

**Supplementary Table 1. Characteristics of mice on the 1g/L N-acetylcysteine water trial ordered by experimental group and date of birth**

| Identification | Sex | Treatment Group | Date of Birth | Treatment Start Date | Age at Euthanasia (wk) | Thymus Weight (g) | Cell Count (x $10^6$) | CD19 Frequency |
|---|---|---|---|---|---|---|---|---|
| 680 | f | NAC | 28-01-21 | 18-02-21 | 16.29 | - | - | - |
| 685 | f | NAC | 28-01-21 | 18-02-21 | 14.00 | - | 80.5 | 99.7 |
| 696 | m | NAC | 28-02-21 | 21-03-21 | 14.14 | - | 58.0 | 99.2 |
| 697 | m | NAC | 28-02-21 | 21-03-21 | 18.71 | - | - | - |
| 511 | f | NAC | 29-03-21 | 19-04-21 | 14.14 | 0.6950 | 180.0 | 99.4 |
| 514 | f | NAC | 29-03-21 | 19-04-21 | 14.14 | 0.9230 | 152.0 | 99.6 |
| 851 | m | NAC | 20-04-21 | 12-05-21 | 20.14 | - | - | - |
| 856 | m | NAC | 20-04-21 | 19-04-21 | 20.14 | - | - | - |
| 537 | f | NAC | 11-05-21 | 02-06-21 | 16.29 | - | - | - |
| 545 | f | NAC | 12-05-21 | 02-06-21 | 22.29 | 0.0430 | 14.5 | 84.8 |
| 546 | f | NAC | 12-05-21 | 02-06-21 | 22.71 | - | - | - |
| 557 | m | NAC | 24-05-21 | 14-06-21 | 16.14 | - | - | - |
| 558 | m | NAC | 24-05-21 | 14-06-21 | 15.14 | - | - | - |
| 962 | f | NAC | 16-06-21 | 07-07-21 | 15.86 | 0.8350 | 222.0 | 99.5 |
| 963 | f | NAC | 16-06-21 | 07-07-21 | 14.14 | - | - | - |
| 690 | f | Control | 19-02-21 | 03-12-21 | 15.43 | - | 167.0 | 99.7 |
| 692 | f | Control | 19-02-21 | 03-12-21 | 10.57 | - | 34.6 | 99.6 |
| 512 | m | Control | 29-03-21 | 19-04-21 | 18.29 | 0.3553 | - | - |
| 513 | m | Control | 29-03-21 | 19-04-21 | 19.00 | - | - | - |
| 853 | f | Control | 20-04-21 | 19-04-21 | 14.00 | 0.9810 | 248.0 | 99.7 |
| 854 | f | Control | 20-04-21 | 19-04-21 | 15.86 | - | - | - |
| 858 | f | Control | 20-04-21 | 19-04-21 | 18.14 | 0.1558 | - | - |
| 552 | m | Control | 14-05-21 | 04-06-21 | 17.57 | - | - | 94.7 |
| 555 | m | Control | 14-05-21 | 04-06-21 | 20.00 | 0.6862 | 120.0 | - |

| | | | | | | | | |
|---|---|---|---|---|---|---|---|---|
| 593 | f | Control | 07-06-21 | 28-06-21 | 11.29 | 0.7637 | - | - |
| 597 | f | Control | 07-06-21 | 28-06-21 | 7.29 | 0.0469 | 78.4 | 27.3 |
| 574 | m | Control | 14-06-21 | 07-07-21 | 18.14 | 0.4606 | 138.0 | 97.4 |
| 575 | m | Control | 14-06-21 | 07-07-21 | 16.14 | 0.3640 | 83.0 | 99.6 |
| 057 | f | Control | 01-08-21 | 23-08-21 | 14.71 | 0.4814 | 210.0 | 99.7 |
| 058 | f | Control | 01-08-21 | 23-08-21 | 17.43 | 0.6123 | 178.0 | 99.5 |

**Supplementary Table 2. Characteristics of mice on the 6.5g/L N-acetylcysteine water trial**

| Identification | Sex | Treatment Group | Date of Birth | Treatment Start Date | Date of Euthanasia | Thymus Weight (g) | Cell Count (x $10^6$) | CD19 Frequency |
|---|---|---|---|---|---|---|---|---|
| 330 | f | NAC | 25-11-21 | 16-12-21 | | | | |
| 333 | f | NAC | 25-11-21 | 16-12-21 | 28-03-22 | 0.7423 | 190.4 | 98.9 |
| 362 | f | NAC | 04-12-21 | 23-12-21 | 28-03-22 | 0.4214 | 207.2 | 99.8 |
| 363 | f | NAC | 04-12-21 | 23-12-21 | 07-04-22 | 0.2487 | 55.7 | 99.1 |
| 435 | f | NAC | 26-01-22 | 17-02-22 | 13-05-22 | - | - | - |
| 436 | f | NAC | 26-01-22 | 17-02-22 | 09-05-22 | - | - | - |
| 437 | m | NAC | 26-01-22 | 17-02-22 | | | | |
| 438 | m | NAC | 26-01-22 | 17-02-22 | 05-05-22 | 0.3254 | 54 | 99.6 |
| 488 | f | NAC | 16-02-22 | 09-03-22 | | | | |
| 489 | f | NAC | 13-02-22 | 09-03-22 | | | | |
| 490 | f | NAC | 16-02-22 | 09-03-22 | 18-05-22 | 0.5324 | - | - |
| 551 | m | NAC | 28-03-22 | 19-04-22 | | | | |
| 553 | m | NAC | 28-03-22 | 03-05-22 | | | | |
| 597 | f | NAC | 11-04-22 | 03-05-22 | | | | |
| 599 | f | NAC | 11-04-22 | 16-12-21 | | | | |
| 358 | f | Control | 02-12-21 | 23-12-21 | 14-04-22 | 0.7874 | 116 | - |
| 359 | f | Control | 02-12-21 | 23-12-21 | 22-03-22 | 1.2824 | 244 | 99.5 |
| 432 | f | Control | 26-01-22 | 17-02-22 | 14-04-22 | 0.7123 | 118 | - |
| 433 | f | Control | 26-01-22 | 17-02-22 | | | | |
| 457 | m | Control | 11-02-22 | 04-03-22 | | | | |
| 458 | m | Control | 11-02-22 | 04-03-22 | | | | |
| 491 | f | Control | 16-02-22 | 09-03-22 | | | | |
| 492 | f | Control | 16-02-22 | 09-03-22 | | | | |
| 494 | f | Control | 16-02-22 | 09-03-22 | | | | |

| | | | | |
|---|---|---|---|---|
| 548 | m | Control | 28-03-22 | 19-04-22 |
| 558 | m | Control | 28-03-22 | 19-04-22 |
| 552 | f | Control | 28-03-22 | 19-04-22 |
| 554 | f | Control | 28-03-22 | 19-04-22 |

**Supplementary Table 3. Characteristics of mice on the 11-week N-acetylcysteine water trial**

| Identification | Sex | Treatment Group | Date of Birth | Treatment Start Date | Date of Euthanasia | Thymus Weight (g) | Cell Count (x $10^6$) | CD19 Frequency |
|---|---|---|---|---|---|---|---|---|
| 051 | m | NAC | 29-07-21 | 19-08-21 | 14-10-21 | 0.0301 | 23.6 | 5.81 |
| 053 | m | NAC | 29-07-21 | 19-08-21 | 14-10-21 | 0.0383 | 32.4 | 12.40 |
| 230 | m | NAC | 18-10-21 | 08-11-21 | 03-01-22 | 0.0542 | 32.0 | 58.80 |
| 232 | m | NAC | 18-10-21 | 08-11-21 | 03-01-22 | 0.0486 | 54.0 | 3.13 |
| 237 | f | NAC | 27-10-21 | 27-10-21 | 12-01-22 | 0.0762 | 113.6 | 2.49 |
| 238 | f | NAC | 27-10-21 | 27-10-21 | 12-01-22 | 0.0705 | 124.8 | 1.37 |
| 301 | f | NAC | 09-11-21 | 09-11-21 | 25-01-22 | 0.0895 | 119.2 | 0.68 |
| 303 | f | NAC | 09-11-21 | 09-11-21 | 25-01-22 | 0.0799 | 136.8 | 1.73 |
| 060 | m | Control | 01-08-21 | 22-08-21 | 19-10-21 | 0.0420 | 10.3 | 88.20 |
| 061 | m | Control | 01-08-21 | 22-08-21 | 19-10-21 | 0.0445 | 6.6 | 82.80 |
| 226 | f | Control | 06-10-21 | 06-10-21 | 22-12-21 | 0.0701 | 107.0 | 5.60 |
| 229 | f | Control | 06-10-21 | 06-10-21 | 22-12-21 | 0.0162 | 52.0 | 0.71 |
| 235 | f | Control | 27-10-21 | 27-10-21 | 12-01-22 | 0.1772 | 30.4 | 91.20 |
| 236 | f | Control | 27-10-21 | 27-10-21 | 12-01-22 | 0.1423 | 24.8 | 89.10 |
| 329 | m | Control | 25-11-21 | 16-12-21 | 10-02-22 | 0.0773 | 21.0 | 41.80 |
| 331 | m | Control | 25-11-21 | 16-12-21 | 10-02-22 | 0.0738 | 88.0 | 0.52 |

**Supplementary Table 4. Characteristics of mice on the littermate 11-week N-acetylcysteine water trial**

| Identification | Sex | Birth Date | Treatment Start Date | Water Group | Date of Euthanasia | Days Alive | Weeks Alive | Thymus Weight (g) | Cell Count (x $10^6$) | CD19 Frequency |
|---|---|---|---|---|---|---|---|---|---|---|
| 852 | f | 29-Jul-22 | | H2O | 11-Nov-22 | 105.00 | 15.00 | 0.2717 | 148 | |
| 856 | f | 29-Jul-22 | | H2O | 11-Nov-22 | 105.00 | 15.00 | 0.2861 | 96 | |
| 961 | f | 18-Sep-22 | | NAC | 05-Dec-22 | 78.00 | 11.14 | 0.052 | 85.6 | 8.09 |
| 968 | f | 18-Sep-22 | | NAC | 05-Dec-22 | 78.00 | 11.14 | 0.0572 | 86.4 | 2.26 |
| 965 | f | 18-Sep-22 | | H2O | 05-Dec-22 | 78.00 | 11.14 | 0.0459 | 43.2 | 22.7 |
| 986 | m | 06-Oct-22 | | NAC | 22-Dec-22 | 77.00 | 11.00 | 0.0805 | 15 | 78.6 |
| 987 | m | 06-Oct-22 | | NAC | 22-Dec-22 | 77.00 | 11.00 | 0.0438 | 20 | 51 |
| 985 | m | 06-Oct-22 | | H2O | 22-Dec-22 | 77.00 | 11.00 | 0.0554 | 3.25 | 85.5 |
| 994 | m | 11-Oct-22 | | H2O | 10-Jan-23 | 91.00 | 13.00 | 0.0621 | 8 | 79.4 |
| 988 | m | 11-Oct-22 | | H2O | 10-Jan-23 | 91.00 | 13.00 | 0.1256 | 17.2 | 89.7 |
| 989 | m | 11-Oct-22 | | H2O | 10-Jan-23 | 91.00 | 13.00 | 0.0799 | 22.8 | 94.5 |
| 995 | m | 11-Oct-22 | | NAC | 10-Jan-23 | 91.00 | 13.00 | 0.0444 | 32 | 4.29 |
| 996 | m | 11-Oct-22 | | NAC | 10-Jan-23 | 91.00 | 13.00 | 0.0287 | 30 | 14.5 |
| 65 | f | 18-Nov-22 | | NAC | 03-Feb-23 | 77.00 | 11.00 | 0.0649 | 21 | 70.6 |
| 66 | f | 18-Nov-22 | | NAC | 03-Feb-23 | 77.00 | 11.00 | 0.0602 | 20.6 | 83.7 |
| 68 | f | 18-Nov-22 | | H2O | 03-Feb-23 | 77.00 | 11.00 | 0.1071 | 26.8 | 90.8 |

| | | | | | | | | | | |
|---|---|---|---|---|---|---|---|---|---|---|
| 69 | f | 18-Nov-22 | | H2O | 03-Feb-23 | 77.00 | 11.00 | 0.1237 | 29.6 | 87.6 |
| 67 | m | 18-Nov-22 | | NAC | 03-Feb-23 | 77.00 | 11.00 | 0.0507 | 7 | 75.4 |
| 70 | m | 18-Nov-22 | | NAC | 03-Feb-23 | 77.00 | 11.00 | 0.0346 | 9.4 | 70.4 |
| 71 | m | 18-Nov-22 | | H2O | 03-Feb-23 | 77.00 | 11.00 | 0.0689 | 19 | 88 |
| 84 | f | 24-Nov-22 | | NAC | 09-Feb-23 | 77.00 | 11.00 | 0.0368 | 64 | 3.9 |
| 85 | f | 24-Nov-22 | | NAC | 09-Feb-23 | 77.00 | 11.00 | 0.0859 | 24.8 | 69.6 |
| 88 | f | 24-Nov-22 | | H2O | 09-Feb-23 | 77.00 | 11.00 | 0.025 | 60.8 | 3.2 |
| 89 | f | 24-Nov-22 | | H2O | 09-Feb-23 | 77.00 | 11.00 | 0.0829 | 49.6 | 31.1 |
| 82 | m | 24-Nov-22 | | NAC | 23-Feb-23 | 91.00 | 13.00 | 0.0249 | 15.4 | |
| 83 | m | 24-Nov-22 | | NAC | 23-Feb-23 | 91.00 | 13.00 | 0.089 | 25.6 | |
| 72 | f | 24-Nov-22 | | H2O | 23-Feb-23 | 91.00 | 13.00 | 0.1119 | 10.6 | |
| 74 | f | 24-Nov-22 | | H2O | 23-Feb-23 | 91.00 | 13.00 | 0.1966 | 18.2 | |
